# *In vitro* efficacy and *in vivo* toxicity and retention of targeted nanoformulated carboplatin in a sustained release carrier for treatment of osteosarcoma

**DOI:** 10.1101/2024.05.02.590788

**Authors:** Kevin Day, Marije Risselada, Marina Sokolsky-Papkov

## Abstract

**Objective:** evaluate 1) if targeting of platinum magnetic nanoclusters will promote uptake in osteosarcoma cells *in vitro*, 2) targeting will improve uptake and delivery in murine OSA *in vivo* compared to free carboplatin, 3) incorporation into a sustained release carrier (SRC) will prolong local retention *in vivo*.

**Methods:** Complex stability and peptide loading was assessed. Drug release was tested at pH 7.4 and 5.5 and cellular uptake and cytotoxity determined for canine, human and mouse osteosarcoma. Subcutaneous murine osteosarcoma was induced and optimal dose and time until tumor growth were established. Tumor bearing mice were equally distributed between 8 treatment (0.5mg carboplatin/mouse) and 1 control group and sacrificed at 8 predetermined time points between 1 hour and 8 days. Blood, tumor site and organs were harvested for tissue ferron and platinum content analysis (ICP-MS).

**Results:** Carboplatin was preferentially released at pH5.5. Targeting increased cellular uptake for carboplatin 15.2-fold, and decreased IC_50_ at 24h and 48h. At 2 weeks, a SC injection of 1-1.5^6^ live cells/mouse reliably resulted in a palpable tumor. Plasma platinum peaked prior to 6 hours while plasma ferron peaked at 24-48 hours. Intratumoral delivery did not lead to a sustained local presence while local delivery in a SRC after surgery did.

**Conclusions:** Targeting of MNC-carboplatin is possible with an increased osteosarcoma cell uptake *in vitro*. *In vivo* metastatic uptake could not be assessed due to lack of metastases, but local delivery in a SRC yielded high local, and low systemic platinum concentrations in mice.

## Introduction

Localized cancer therapy has attracted considerable attention due to the ability to deliver higher local drug load, regardless of vascular status, and reduce systemic toxicity, with various delivery systems used (such as hydrogels,^Risselada-2017a, Risselada2016,Risselada 2020^ polymers^Vishwarao2016^ and calcium sulfate hemihydrate^Tulipan2016, Tulipan2017,Maxwell2020,Phillips2018,Risselada2015,Worth2020, Belda2021^). In addition, local drug delivery systems have been used to visualize and/or treat metastasis in a theranostic approach.^Li2012^ Magnetic nanoparticles (MNPs) have long been studied as magnetic resonance (MR) imaging agents and are known for their biocompatibility and low toxicity.^Singh2014a, Singh2014b^ Their presence could allow assessment of uptake in residual disease as well as distribution into metastatic lesions. The authors previously developed a simple theranostic nanoformulation based on magnetic nanoparticles stabilized by a bisphosphonate-modified poly(glutamic acid)-b-(ethylene glycol) block copolymer (MNC) and complexed with platinum drugs.^Vishwarao2016^ Targeting the formulation with peptides based on cell specific surface receptors would allow selective delivery of carboplatin into cells. Two potential cell surface receptors have been identified for osteosarcoma (OSA): Insulin-like Growth Factor-1 Receptor (IGF-1R) described in both canine and human OSA^Hassan2012,Friebele2015, Schmidt2013,Rodriguez2014,Schiffman2015,Maniscalco2015,^ and Ephrin type-A receptor-A (EphA-2),^Posthuma2016^ a recently discovered receptor in human OSA, but not described in canine OSA yet. Prior work showed that targeting EphA-2 receptors in ovarian cancer cells was possible.^Scarberry2008^ Biodegradable or thermo-sensitive injectable or implantable polymers and gels can be used for sustained local delivery of drugs.^Sokolsky-Papkov2009, Mathews2009,Golovanevski2015, Risselada2016,Risselada2017a, Risselada2020,Risselada2024^ Polymers based on polylactic acid-castor oil are solid to viscous liquid in RT based on their composition, biocompatible and were shown to incorporate and release bupivacaine *in vivo*, prolonging the local analgesia to 96 hours,^Sokolsky-Papkov2009,Golovanevski2015^ and release rates can be tuned further based on the polymer composition. These polymers can successfully incorporate and release nanoparticles.^Lee2017^ Tissue adherent microgels based on oxidized dextran form an in situ-depot by reacting with amino groups present in the proteins in the injection site.^Denga2016^ We propose to incorporate targeted MNC formulated carboplatin (t-MNC-carboplatin) into injectable polymers based on polylactic acid-castor oil and tissue adherent microgels.

The aims for this study were to: 1) evaluate if t-MNC-carboplatin can selectively promote the uptake of carboplatin into cancer cells and enhance its activity against murine OSA *in vitro*, 2) t-MNC-carboplatin will improve uptake and delivery in murine OSA *in vivo* compared to free carboplatin, 3) incorporation into a sustained release carrier (SRC) will prolong local retention *in vivo*. Our hypotheses were that 1) t-MNC-carboplatin would be stable *in vitro* with a similar IC_50_ and an increased cellular uptake compared to free carboplatin and non-targeted MNC-carboplatin, 2) targeting would increase uptake *in vivo*, and would lead to an improved outcome compared to free carboplatin while 3) incorporation into a SRC would prolong local retention.

## Materials and Methods

### *MNCs preparation, in vitro* stability, IC50 and cellular uptake

#### Particle loading and *in vitro* drug release

Magnetic nanoclusters (MNCs) were prepared as previously described.^Vishwarao2016^ Specifically, magnetic nanoparticles (MNPs) with an average diameter of 9nm were stabilized by a bisphosphonate-modified by polyglutamic acid homopolymer or poly(glutamic acid)-b-(ethylene glycol), poly(aspartic acid)-b-(ethylene glycol) block copolymers. Cisplatin or carboplatin was loaded as previously described and the drug loading was measured by ICP-MS.^Vishwarao2016^ MNCs stability with or without drug:

The targeting peptides based on EphA-2 (EphA-2-binding peptide, peptide 66) and IGF-1R (IGF-1R-binding peptide, peptide 67) were conjugated to the polymer carboxylic side chains through EDC/S-NHS chemistry and purified using filter centrifugation. The conjugation efficiency and the targeting peptide content was determined by HPLC analysis of non-conjugated peptide. The polylactic acid-castor oil block copolymer for the SRC was prepared as previously described^Sokolsky-Papkov2009b^ at 50:50 ratio of castor oil to lactic acid. The sustained release formulation was prepared by directly mixing the carboplatin/t-MNC-carboplatin into the polymer at 5% w/w. The particles were dispersed in PBS pH 7.4 and ABS pH 5.5 and the Pt release was measured by ICP-MS as previously described.^Vishwarao2016^

A 14-day elution curve was obtained of 1) t-MNC-carboplatin, 2) carboplatin in SRC x 3; 3) t-MNC-carboplatin in SRC. The release of the drug-loaded nanoparticles was compared to the release of the free drug. The load % of coated particles was measured using ICP-MS as previously described.^Vishwarao2016^

#### *In vitro* cellular uptake and cytotoxicity

Enhancement of cellular uptake by OSA cells by using the proposed targeting peptides was evaluated as follows: 6-well plates were seeded with 10^6^ cells each and treated for 24hrs with: 1) carboplatin; 2) MNC-carboplatin and 3) t-MNC-carboplatin, after which the cells were washed, harvested and lysed. Pt and Fe cellular content was measured by ICP-MS using a prior described protocol.^Vishwarao2016^

*In vitro* cytotoxicity of the new carboplatin formulation was assessed in canine OSA: OSCA-40 (cell line made available for *in vitro* use by Dr M. Hauck) and D17 (ATCC, Manassas, Virginia, US) as well as human OSA (CRL-1543, ATCC) and mouse OSA (K7M2wt, ATCC) for carboplatin and cisplatin. Twenty-four hours after seeding about 3000 cells/well in 96-well plates, the cells were treated for 24 and 48 hrs. A colorimetric assay (Cell Titer Blue Cell Viability Assay, Promega, Madison, WI, US) that detects cellular metabolic activities was used to assess cell viability. The cell viability was determined by comparing the different treatment groups with the control (untreated) wells. Three different treatment combinations were evaluated for each cell line: 1) carboplatin; 2) MNC-carboplatin; 3) t-MNC-carboplatin; 4) cisplatin; 5) MNC-cisplatin and 6) t-MNC-cisplatin.

### *In vivo* murine subcutaneous osteosarcoma model

Institutional IACUC approval was obtained (PACUC #1711001644) for this study. Osteosarcoma cells (K7M2wt, ATCC) were injected in the subcutaneous space over the dorsum of fourteen female BALB/c mice of approximately 20-gram body weight (Charles River). Live cells were harvested, diluted and suspended in 100uL ice-cold sterile buffered saline per dose, according to prior published protocols ^Overwijk2001^ Three doses were evaluated for tumor growth: 2^5^ (n=4) 5^5^ (n=4) and 1^6^ (n=6). Mice were monitored daily to pinpoint a day post inoculation at which tumor growth reliably occurred to use for the remainder of the study.

### *In vivo* outcome, local retention, distribution and efficacy

*In vivo* efficacy was evaluated using the optimized mouse model: 1.5^6^ K7M2wt cells in 100uL were injected subcutaneously over the dorsum of 192 female BALB/c mice. Eight treatment groups were included, and three mice were sacrificed at 8 time points [1 hour, 6 hours, 12 hours, 1 day, 2 days, 3 days, 5 days, and 8 days] for each group. Treatment groups were: control (control), intraperitoneal (IP) non-targeted MNC-carboplatin with surgery to remove the primary tumor; IP non-targeted MNC-carboplatin without surgery to remove the primary tumor; IP t-MNC-carboplatin with surgery to remove the primary tumor; IP t-MNC-carboplatin without surgery to remove the primary tumor; intratumoral injection of t-MNC-carboplatin without surgery; surgical removal of the primary with local delivery of t-MNC-carboplatin in polymer; surgical removal of the primary with local delivery of non-targeted MNC-carboplatin in polymer. The total dose per mouse was 0.5mg carboplatin in a total volume of 0.1ml. The drug was filtered immediately prior to administration (20micron filter, Corning; Kentucky, USA) and aseptically handled thereafter. The formulation in a SRC was aseptically handled but not filtered immediately prior to administration. Mice were checked daily for incisional complications and overall health. After euthanasia, the wound bed and all internal organs were grossly examined.

The wound bed was evaluated for necrosis, dehiscence and for residual or recurrent disease. The ventral and dorsal aspects of the lungs were photographed to allow quantification of any metastatic lesions present within the lung and the number/size of metastases were counted and measured. Samples obtained were: plasma, primary tumor/wound bed, lungs (areas macroscopically free and with macroscopic metastases were collected separately) as well as liver and kidneys. All samples were weighed prior to freezing to −70°C and were processed in one batch for measurement of Pt & Fe content using ICP-MS analysis.

## Results

### *In vitro* evaluation of stability, IC50 and cellular uptake

#### Particle loading and *in vitro* drug release

Particle loading (expressed as drug load %) ranged from 5.12-9.24% (Table 1, Table 2, Fig 1) with load % of the targeted formulation lower but comparable, and drug load of cisplatin higher than carboplatin. All particles released fully from the polymer, with an increased drug release in pH 7.4 and when coupled to a peptide (Figure 1). Drug release was increased in pH 7.4 compared to pH 5.5 and when coupled to a peptide. Cisplatin did not exhibit the same pH triggered release post peptide conjugation while carboplatin release increased post peptide conjugation (Figure 2).

**Figure 1:**
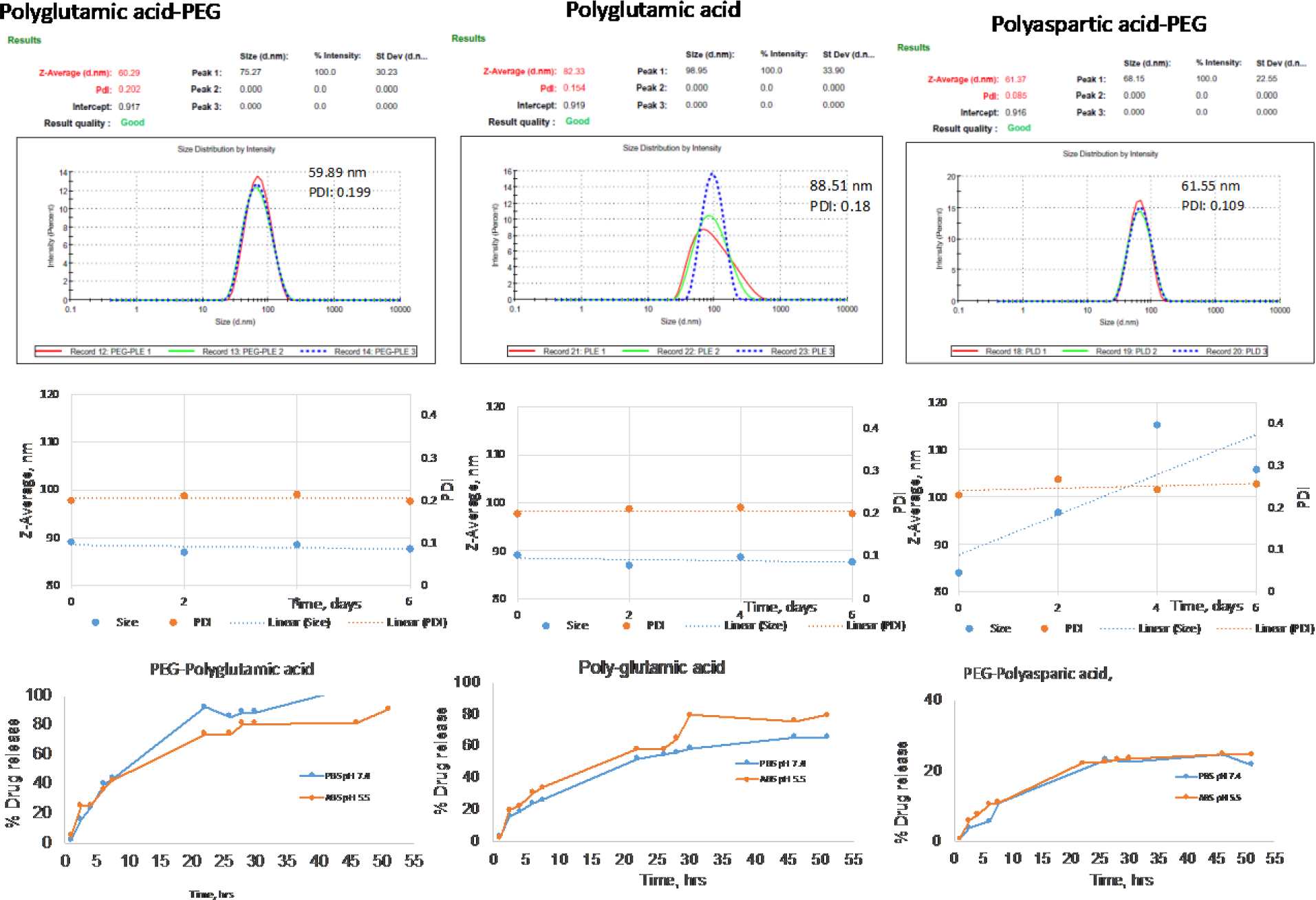
Summary of *in vitro* assessment of nanoparticles pre-targeting. The particles size, PDI, stability of the carboplatin loaded particles and the pH triggered release were assessed. Polyglutamic coated particles were more stable and exhibited a greater pH release effect.

**Figure 2:**
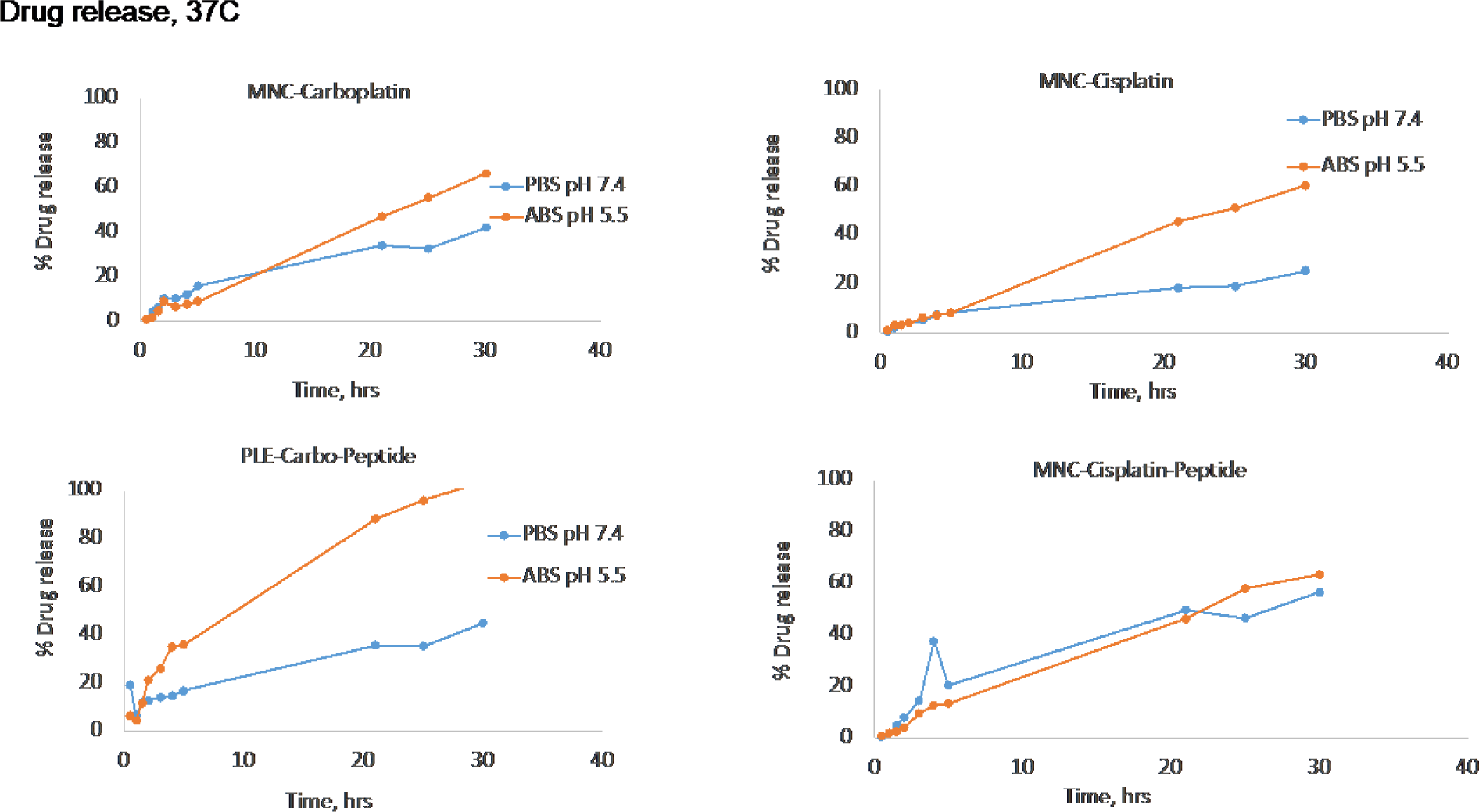
Percentage drug release *in vitro*. Release of MNC-carboplatin (A) with or without coupling to a peptide (B) was compared for an environment with a pH of 7.4 and 5.5. Drug release was increased in pH 7.4 compared to pH 5.5 and when coupled to a peptide. Cisplatin did not exhibit the same pH triggered release post peptide conjugation while carboplatin release increased post peptide conjugation.

**Table 1.** Drug load percentage (measured using ICP-MS) expressed as %.

| Sample | Drug load% |
| --- | --- |
| MNC-carboplatin | 6.26 |
| MNC-carboplatin-peptide | 5.12 |
| MNC-cisplatin | 9.24 |
| MNC-cisplatin-peptide | 8.20 |

Drug loading for both targeted and non-targeted carboplatin and cisplatin is shown. MNC=magnetic nanoclusters

**Table 2.** Influence of coating on drug loading (measured using ICP-MS) expressed as %. Polyglutamic coated particles had a higher carboplatin load %.

| Sample | Fe % | Fe<sub>3</sub>O<sub>4</sub> % | Carboplatin load % |
| --- | --- | --- | --- |
| PEG113-PLD50 (PEG-Aspartic) | 13.38 | 18.46 | 9.24 |
| PLE 100 (Glutamic acid) | 26.50 | 36.57 | 7.74 |
| PEG113-PLE100 (PEG-Glutamic) | 21.40 | 29.53 | 15.19 |

#### *In vitro* cellular uptake and cytotoxicity

Conjugating with a peptide used to target EphA-2 (peptide 66) more effectively increased cellular uptake compared to conjugating with an IGF-1R-binding peptide (peptide 67) or to non-conjugated PLE (Table 3, Figure 3). This effect was seen for all 4 cell lines. The % cellular uptake in mouse OSA cells (K7M2wt) was higher for free cisplatin and MNC loaded cisplatin compared to their respective carboplatin formulations (Figure 4). MNC loading did not influence cellular uptake. While targeting of the MNC-formulation increased the cellular uptake for both cisplatin and carboplatin, this increase was more pronounced for carboplatin and the cellular uptake for targeted-MNC-carboplatin was the highest for all formulations assessed. The MNC formulation decreased the cytoxicity, while conjugating with a peptide to create a targeted formulation increased the cytotoxicity (Table 4). Ultimately carboplatin conjugated with EphA-2 as targeting peptide was chosen for the *in vivo* part of the study due to better uptake in the cells and a higher differential between targeted versus non-targeted formulations.

**Figure 3:**
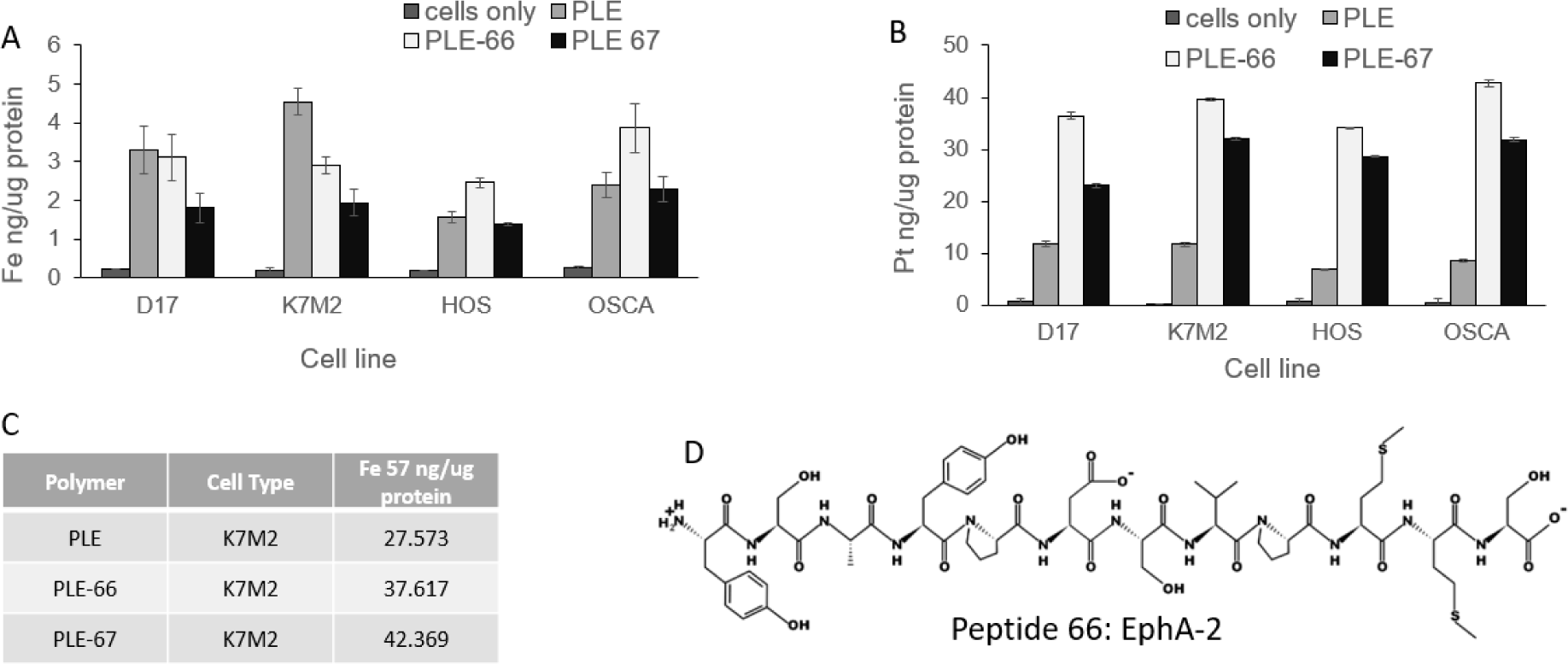
Assessment of two targeting peptides. A, B uptake of Ferron and Platinum in 4 osteosarcoma cell lines. C: uptake in murine osteosarcoma, D: Structure of peptide 66. Peptide 66 was targeted at binding to EphA-2, peptide 67 targeted at IGF-1R. Conjugating with EphA-2-binding peptide resulted in increased cellular uptake for murine (K7M2wt), human (HOS) and canine (OSCA) osteosarcoma cells *in vitro* and was chosen as the targeting peptide for the remainder of the study.

**Figure 4.**
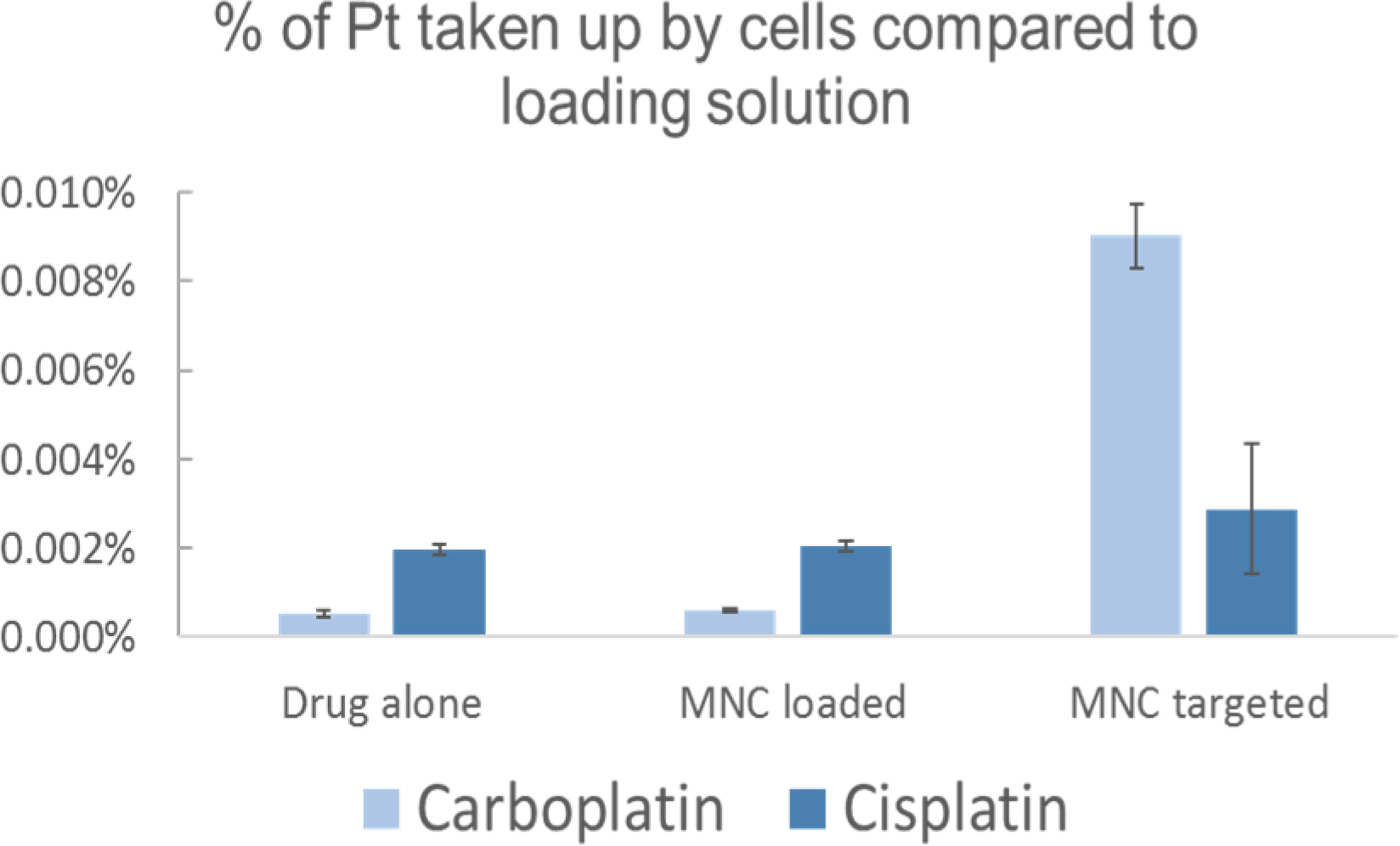
Cellular uptake of platinum in K7M2wt *in vitro*. Platinum uptake in cells for carboplatin and cisplatin in an (EphA-2-) targeted and non-targeted MNC formulation compared to non-altered formulation. Targeting improved cellular uptake, both in cisplatin and carboplatin.

**Table 3.** Drug load percentage of non-targeted and targeted coated particles (measured using ICP-MS) is expressed as %).

| Sample | Carboplatin load %, |
| --- | --- |
| PLE carboplatin | 1.36 |
| PLE 66 (EphA-2 targeted) carboplatin | 2.98 |
| PLE 67 (IGF-1R targeted) carboplatin | 2.66 |

**Table 4.** IC_50_ (after 24h hours incubation) determination of non-targeted clusters: no significant differences between formulations. Polyglutamic coated particles performed better than polysaspartic coated particles. HOS: Human osteosarcoma, K7M2wt: murine osteosarcoma, D17 and OSCA: canine osteosarcoma.

| Cell type | Drug and Coating | IC<sub>50</sub> ug/ml |
| --- | --- | --- |
| HOS | **Carboplatin** | **22.09** |
|  | Polyglutamic-PEG (PLE-PEG) | 29.15 |
|  | Polyglutamic acid (PLE) | 28.31 |
|  | Polyaspartic acid-PEG (PLD-PEG) | 28.28 |
| K7M2wt | **Carboplatin** | **18.39** |
|  | Polyglutamic-PEG (PLE-PEG) | 26.98 |
|  | Polyglutamic acid (PLE) | 15.35 |
|  | Polyaspartic acid-PEG (PLD-PEG) | 24.38 |
| D17 | **Carboplatin** | **119.89** |
|  | Polyglutamic-PEG (PLE-PEG) | 99.83 |
|  | Polyglutamic acid (PLE) | 103.77 |
|  | Polyaspartic acid-PEG (PLD-PEG) | 142.93 |
| OSCA | **Carboplatin** | **117.59** |
|  | Polyglutamic-PEG (PLE-PEG) | 80.39 |
|  | Polyglutamic acid (PLE) | 81.98 |
|  | Polyaspartic acid-PEG (PLD-PEG) | 83.17 |

### *In vivo* murine osteosarcoma model

None of the mice inoculated with 2^5^ or 5^5^ cells grew macroscopically visible or palpable tumors at the injection site. Three out of 6 mice inoculated with 1^6^ cells showed palpable tumor growth at 2 weeks. An inoculation dose of 1-1.5^6^ live cells/mouse was chosen for the remainder of the study.

### *In vivo* outcome, local retention, distribution and efficacy

All 192 mice were inoculated with 1-1.5^6^ live K7M2wt cells, and tumors allowed to grow for a minimum of 2 weeks prior to treatment at d=0. Total dose per mouse was 0.5mg of carboplatin, or 0.1ml of product. Tumors reliably grew in 128 mice and two mice per time point [1-hour, 6-hour, 12-hour, 1-day, 2-day, 3-day, 5-day, and 8-day groups] were included for all 8 treatment groups.

#### Outcome

Two mice died: one 6-hour mouse [IP t-MNC-carboplatin with surgery] and one 3-day mouse [IP t-MNC-carboplatin w surgery]). Two sites dehisced: the same 3-day mouse [IP t-MNC-carboplatin with surgery] that died and one 1-day mouse [IP non targeted MNC-carboplatin with surgery]. No other incisional or systemic complications were noted.

#### Local retention

No increased local concentration of Pt was found in the control mice, after IP injection of non-targeted MNC carboplatin with or without surgery and after IP injection of targeted MNC carboplatin with or without surgery (Figure 5). High levels of both Pt and Fe were found in the site after intratumoral injection for 12 hours. Local delivery of MNC-carboplatin in a SRC after surgery led to an increase of both Pt and Fe measured at the site with a peak at 120 hours. The non-targeted MNC-carboplatin delivered locally had elevated Pt in the site initially, but with an immediate drop and without a steady increase over time. Platinum and Fe for the local sustained release and intratumoral delivery groups mirrored each other over time (Figure 5).

**Figure 5.**
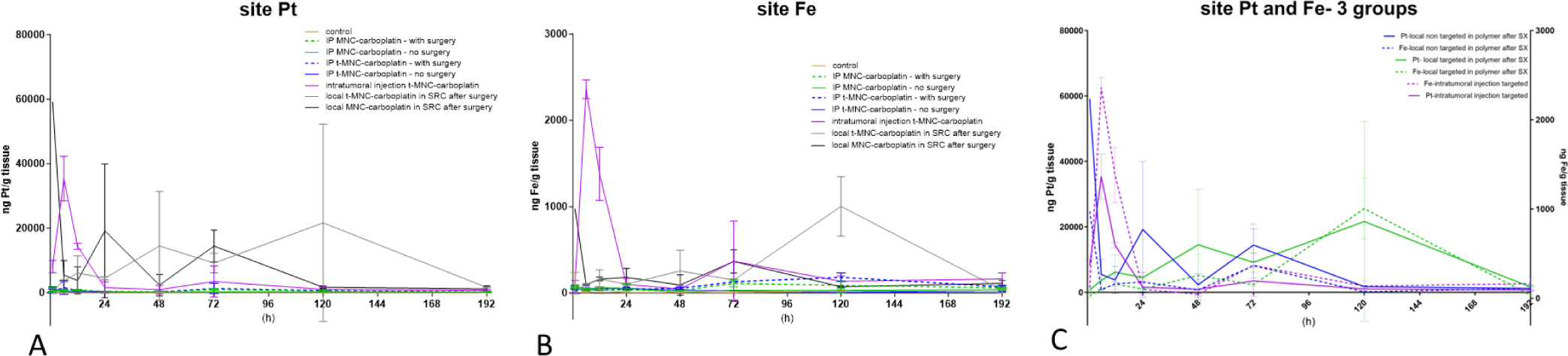
Tumor and surgery site Pt content for all groups (A), Fe content for all groups (B), Pt and Fe content for locally delivered drugs (C). Intratumoral delivery did not lead to a sustained local presence while local delivery after surgery in a sustained release carrier (SRC; polylactic acid-castor oil block copolymer) did.

#### Distribution

Plasma Pt levels for all treated mice were highest within the first 12 hours, with a peak prior to 6 hours and decreased thereafter. Plasma Fe levels showed a later peak at 48 hours with one outlier at 24 hours (Figure 6). No increased Pt was noted in the control group mice in either liver or renal tissue assessed (Figure 7). All mice with IP delivered non-targeted MNC-carboplatin showed high Pt content in renal tissue, whereas the Fe concentration in renal tissue was lower than in lung, and especially liver. There was no high uptake in the surgery site or in non-removed tumors, although uptake seemed slightly higher in sites that had tumors removed versus those that did not. Intraperitoneal t-MNC-carboplatin similarly had high renal Pt content than in liver or lung, but lower Fe renal content, although the effect for Pt was less pronounced. These increases were not seen for locally delivered carboplatin.

**Figure 6.**
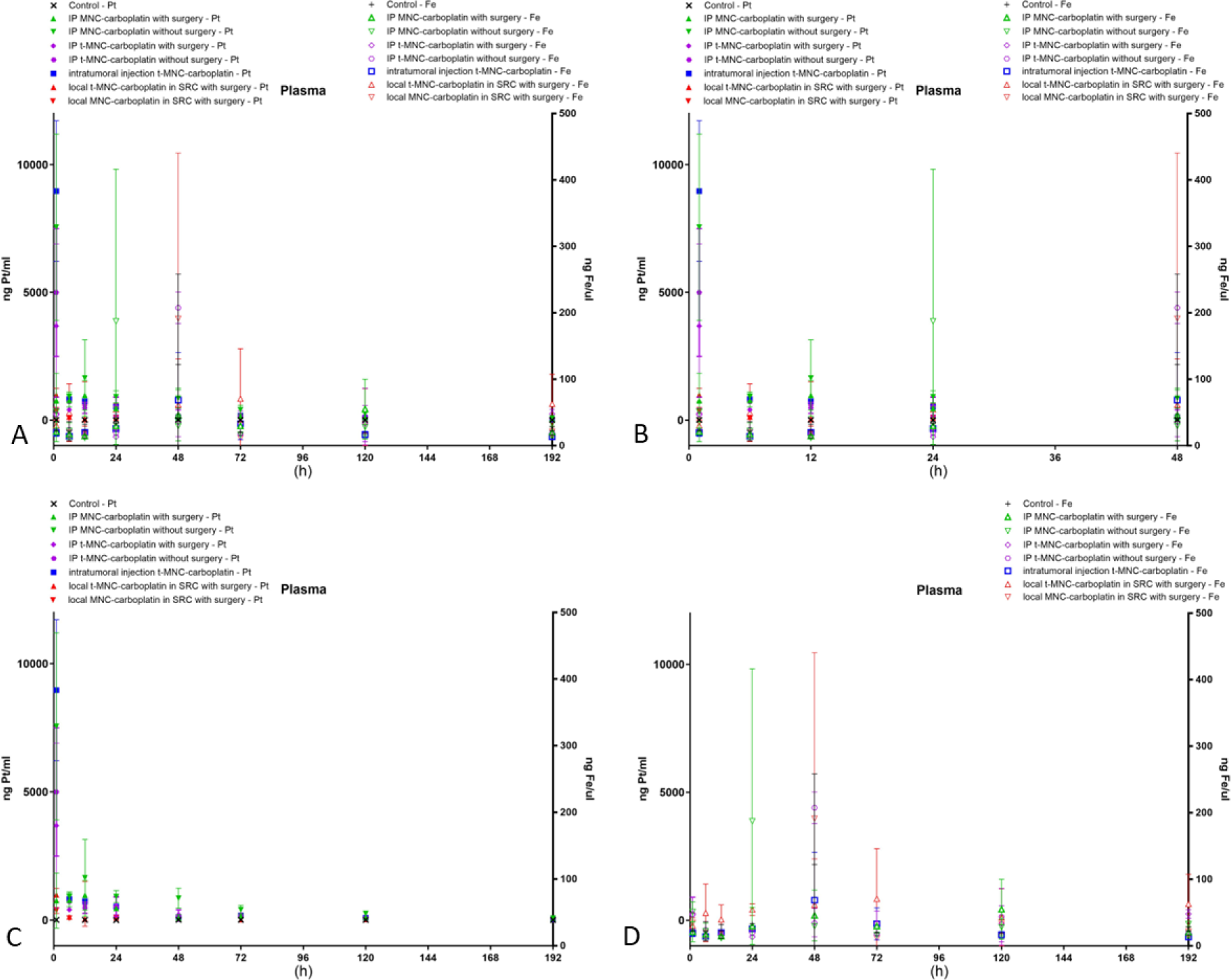
Plasma Platinum (Pt) and Ferron (Fe) are shown for the entire study duration, and for the first 48hours. A: Plasma Pt and Fe over 192 hours, B: Plasma Pt and Fe for the first 48 hours, C: Plasma Pt only over 192 hours; D: Plasma Fe only over 196 hours. Plasma Pt peaked earlier than plasma Fe: Plasma Pt peaked prior to 6 hours whereas plasma Fe peaked at 24-48 hours.

**Figure 7.**
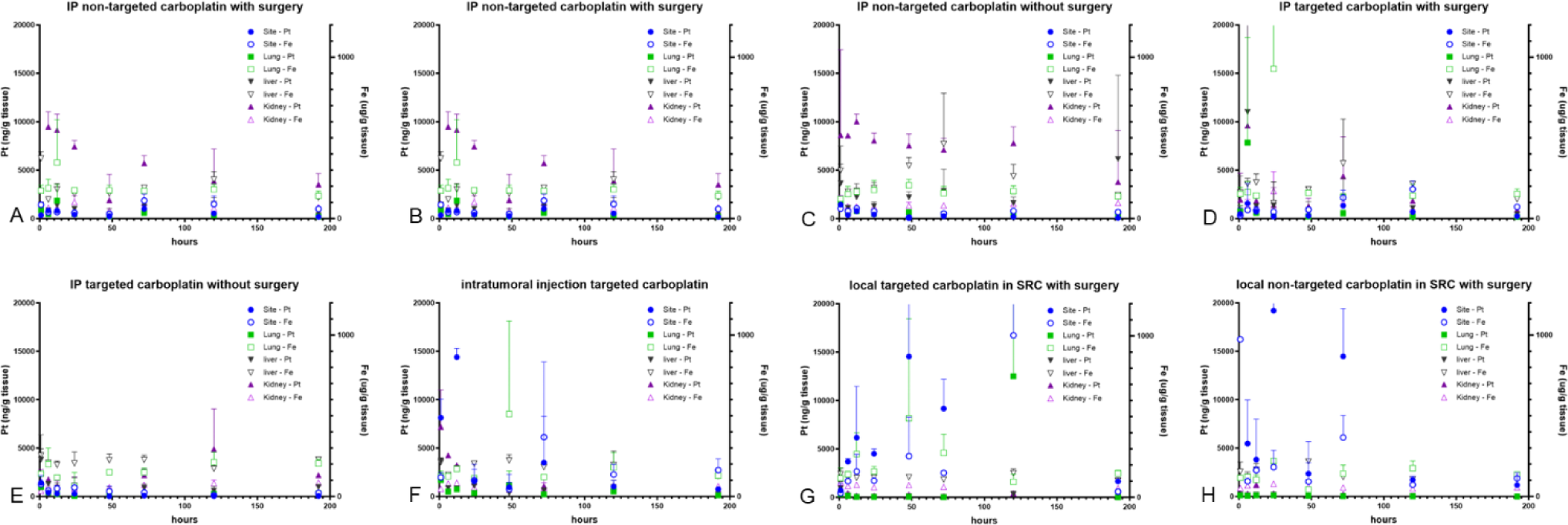
Platinum (Pt) and Ferron (Fe) for implantation site, lung, liver and kidney at time point are shown for the control and 7 treatment groups. For each variable the solid symbol represents Pt, and the corresponding nonsolid symbol/color represents Fe. The left Y-axis shows Pt and the right Y-axis Fe concentration. Locally delivered carboplatin consistently had higher local concentrations with less systemic uptake, whereas IP delivered carboplatin had higher systemic uptake.

#### Efficacy

No macroscopically visible lung metastases were seen in any of the mice. Intraabdominal lesions were seen in 2 mice (one 5-day mouse with disseminated lesions [IP MNC-carboplatin without surgery] and one 6-hour mouse in which the primary tumor grew into the abdomen [control group].

## Discussion

In the *in vitro* portion of this study, polyglutamic coated particles were found to be more stable and exhibited a greater pH release effect than other formulations. Targeting of both nanoformulated carboplatin and cisplatin with either an EphA-2-binding peptide or IGF-1R-binding peptide was possible, and the formulations yielded a similar platinum load as the non-targeted formulations. Targeting increased cellular uptake of the formulation *in vitro* in osteosarcoma cells in across species, with use of an EphA-2-binding peptide providing a more consistent and reliable increase in uptake than non-targeted or use of an IFG-1R-binding peptide. The % uptake by targeting the formulation was more pronounced for carboplatin than for cisplatin. Polyglutamic coated particles & carboplatin targeted with an EphA-2-binding peptide were chosen for the *in vivo* part of the study. A local tumor model with K7M2wt was successfully reproduced in mice but did not create distant metastases. Intratumoral delivery reached high local concentrations initially, but rapidly declined thereafter, without systemic uptake in mice. Local delivery in a sustained release compound yielded high local, and low systemic Pt concentrations. Platinum retention and distribution did not fully mirror Ferron distribution.

Two cell surface receptors were chosen as possible targets to investigate in this study. These were IFG-1R (described in both human and canine osteosarcoma)^Hassan2012,Friebele2015,Schmidt2013,Rodriguez2014,Schiffman2015,Maniscalco2015^ and EphA-2 (described in human but not in canine osteosarcoma).^Posthuma2016^ Neither receptor has been described in mouse osteosarcoma. The presence of EphA-2 has been associated with malignancy in various human cancers.^Wykosky2005,Xiao2020^ The synthetic peptide analogues for both cell surface receptors were successfully incorporated into the targeted formulations investigated in this study, with a platinum load % that was similar, but smaller compared to the non-targeted formulations (5.1% vs 6.3% for carboplatin and 8.2 vs 9.2% for cisplatin).

While the MNC formulation in itself had decreased cellular uptake, targeting of the formulation subsequently increased uptake in OSA cells *in vitro* across species. Targeting based on an EphA-2-surface receptor outperformed targeting based on IFG-1R *in vitro*. While cisplatin had a higher platinum load %, the pH triggered release and increased cellular uptake of carboplatin was why ultimately carboplatin was chosen for the *in vivo* portion of the study. We further chose to include various treatment groups while focusing on one binding peptide (based on an EphA-2-binding peptide).

A subcutaneous osteosarcoma tumor model was recreated using an adaptation of protocols that utilized an injection into an appendicular bone (femur ^Miretti2008^ or proximal tibia ^Cole2011,Crasto2018^). Two mice had a local wound dehiscence, however, this most likely could be attributed to residual tumor in the surgery site rather than secondary to local drug delivery as both mice received carboplatin IP and not locally. The overall lack of local tissue and wound complications was in line with prior studies where carboplatin was administered subcutaneously with a different SRC, either in poloxamer^Risselada2017a^ or in CaSO_4_ beads.^Belda2021^ However, due to differences in species (rats^Risselada2017a, Belda2021^ vs mice) and methodology (local Pt only measured at d7^Risselada2017a^, d28 ^Belda2021^ vs multiple time points up to d7), no direct comparison of the SRC’s ability to local retain carboplatin could be made.

Intratumoral delivery reached high local concentrations initially, but rapidly declined thereafter, without systemic uptake. Local delivery of both targeted and non-targeted MNC-carboplatin in a sustained release compound (polylactic acid-castor oil block copolymer) did yield high local, and low systemic Pt concentrations in mice. This is similar to other local delivery methods.^Belda2021,Risselada2017a^

Targeting of the formulation did not result in peripheral tumor uptake when delivered intraperitoneally, which could be due to the amount of drug taken up in circulation and therefore reaching the tumor site. Local delivery, however, might lead to accumulation in draining (sentinel) lymph nodes via lymphatic drainage, rather than relying on systemic circulation. This effect could be increased if the drug is selectively taken up in osteosarcoma cells due to targeting of the formulation. Further studies utilizing local drug delivery in an area with clear established locoregional draining lymph nodes in a larger sized species might be needed to fully investigate this potential. Platinum retention and distribution did not fully mirror Ferron in the current study. The ability to use the formulation to serve as a theranostic drug and have diagnostic purposes in addition to therapeutic effects, both should have a similar pattern. Follow up studies are therefore needed to fully investigate if Fe uptake in local lymph nodes or metastatic lesions or does mirror each other and therefore would provide the diagnostic arm of the formulation.

### Limitations

Several limitations to the study exist, firstly only low numbers per group were included, and had variation between the values obtained. This precluded making a full pharmacokinetic analysis, and results are therefore more descriptive and observational, and should be seen as pilot data. We purposefully opted to increase the number of variations rather than numbers per group in this pilot project to gather data for future larger studies. Despite the low numbers some conclusions regarding local retention and absence of local toxicity could be drawn. A second limitation was the lack of visible metastases, disallowing the assessment of uptake in metastatic lesions.

Development of pulmonary metastases in a mouse osteosarcoma model has been described prior, both with the same cell line^Crasto2018^, and with different tumor cell lines.^Cole2011,Miretti2008^ It is possible that the chosen injection site in our study (subcutaneous) does not lend itself well to cancer spread compared to an intrafemoral ^Miretti2008^ or proximal tibial^Cole2011,Crasto2018^ injection site. We chose the subcutaneous site, as tumor growth needed to be readily visible, and tumor removal feasible with a minor surgery.

A third limitation was due to the maximum volume that could be delivered locally or intratumorally. This limited the dose that could be delivered, including for IP delivery, as the dose was kept consistent between mice.

### Conclusions

Targeting of MNC-carboplatin was possible with use of EphA-2-binding peptide resulting in an increased osteosarcoma cell uptake *in vitro* for various species. *In vivo* metastatic uptake in mice and efficacy could not be assessed due to lack of metastases, but local delivery in a SRC yielded high local, and low systemic Pt concentrations.

